# Low-abundance taxa drive host microbiome response to temperature stress

**DOI:** 10.1101/2025.09.18.677205

**Authors:** Kaden Muffett, Stephen Williams, Sophia MacVittie, Mara JW Schwiesow, Susan Arechiga Mendoza, Salma Arechiga Mendoza, Hailey Hatch-West, E. Maggie Sogin

## Abstract

Global climate change (GCC) will lead to the deterioration of host and microbiome health, posing a significant threat to complex symbioses. To gauge the reproducibility of thermal warming associated with GCC and the time scales on which it occurs, we monitored shifts in the *Exaiptasia* microbiome in 24 populations of the model cnidarian *Exaiptasia diaphana* over the course of six months. We identified a small cohort of temperature-sensitive amplicon sequence variants (ASVs) that reproducibly declined with increases in temperature before changes in host fitness occurred. After comparing these ASVs to global datasets, we found that the heat-impacted ASVs from our experiment were also present in many other cnidarian species, highlighting their potential as widespread indicators of thermal stress. Overall, we find that fewer than 1% of the microbiome within *E. diaphana* are heat indicator taxa, suggesting that a small but consistent group of ASVs signals the early stages of thermal stress.

## INTRODUCTION

Climate change poses a significant threat to coastal marine ecosystems. As temperatures rise, microbe-dependent marine life, including corals and sponges, will face both host system and microbiome destabilization^1–4^. For cnidarians, increased temperatures disrupt the animal host by increasing metabolism and decreasing tissue integrity^5,6^. For their core algal endosymbionts, heat stress leads to decreased photosynthetic efficiency, increased metabolic demands, and, at the highest temperatures, mortality^3,7,8^. In addition to host and algal destabilization, many marine microbes have distinct thermal maxima^9–11^.

Marine heat waves and ocean temperature significantly impact the composition of complex microbial communities^9,12,13^. As these communities change, so does the function of the assemblage, presenting an additional challenge for host metazoans that rely on a stable microbiome^9^. In microbial communities associated with animal and plant hosts, exposure to heat often creates or exacerbates dysbiosis. Microbiome disruption negatively impacts an impressive suite of functions, such as reproduction, growth, food uptake, and tissue defense^14–18^. In photosymbiotic cnidarians, the bacteriome may facilitate the transfer of lipids and sugars across the symbiosome barrier^19,20^. Destabilization in these systems may cause a breakdown in the nutrient cycling throughout the host, leading to stress^21–23^.

For anthozoans, the microbial communities in lab and field populations are highly complex, bearing little similarity along lines of host phylogeny and showing low continuity across time^20,24–26^. These bacterial community trends across time are replicated within the laboratory model *Exaiptasia diaphana*^27–29^. While the microbiome is known to support the adaptability of anthozoan hosts, these highly diverse and plastic systems make the identification of specific heat-tolerance and heat-stress-associated bacteria difficult^30,31^.

With microbiome work that occurs only after phenotypic destabilization (e.g., bleaching) in coral, we cannot gain insight into early microbial indicators of oncoming thermal stress^32,33^. To observe these dynamics, we used two host-algal pairings: one thermally tolerant (*Exaiptasia* CC7 with *Symbiodinium microadriaticum*) and one thermally sensitive (*Exaiptasia* H2 with *Breviolum pseudominutum*). Using these anemone lines, we tracked the onset of bacteriome change and host distress across three temperatures, ranging from temperate (26.5 °C) to hot (31.5 °C), over six months, and paired these data with intensive sampling during acute stress at an extreme temperature (34 °C).

## RESULTS

To quantify the impact of elevated temperatures on *E. diaphana* performance, we compared anemone population dynamics and host-algal physiology across temperatures and *Exaiptasia* pairings (H2 and CC7). We exposed replicate populations of H2 and CC7 host-algal pairings to ecologically relevant elevated temperatures (+2.5 °C to +7.5 °C) as predicted for climate change scenarios for 2100^34^. As a result of this, we leveraged our thermal stress experiments to explore host-microbiome responses to chronic, long-term exposure to +2.5 °C and +5 °C and acute, short-term exposure to +7.5 °C (34 °C). Our experimental framework enabled us to compare the population-level and physiological responses of *E. diaphana* to elevated thermal conditions across time, while monitoring associated changes in microbiome communities. In doing so, we were able to identify four microbial taxa that have (1) a conserved response across host-algal pairings, (2) are implicated in significant declines in the relative abundance of other microbial taxa, and (3) occur in a range of cnidarian families and genera.

Our population and physiology data revealed that chronic exposure to +5°C negatively impacted H2, but not CC7. To identify the broad microbiome phenotypes of heat exposure in microbiomes, we calculated alpha and beta diversity for our 493 samples across 9 time points in H2 and CC7 at temperatures from baseline to +7.5 °C. To capture specific microbes impacted by temperature exposure, we produced linear models of the core and common microbiome of *E. diaphana* across the 180 days of chronic heat exposure and 14 days of acute heat exposure, then isolated ASVs with a conserved chronic and acute phenotype. Finally, we placed these specific taxa within the broader context of global marine microbiomes.

### Elevated temperatures differentially impact *Exaiptasia* performance and population structure between host-algal pairings

Chronic exposure to elevated temperatures increased anemone reproduction and population growth in CC7 but not H2 host-algal pairings. While CC7 populations grew faster at higher temperatures compared to populations held at baseline (ANOVA; *p =* 0.02; Supplementary Tables S1 and S2), H2 population growth significantly declined at elevated temperatures over time (ANOVA, *p* = 0.0002; Supplementary Tables S2 and S3). Using a generalized additive model (GAM), we fit nonlinear effects to weekly counts of asexual recruits to resolve population changes across temperatures. Our data showed clear non-linear impacts of increased temperature on CC7 and H2 asexual reproduction (Fig. 2b). H2 and CC7 individuals have their highest production of asexual recruits at estimated thermal maxima of 28.5 °C (+2 °C) and 29.5 °C (+3 °C), respectively (ANOVA GAM, *p < 0*.*00001*; Supplementary Table S4). Additionally, the data underlying these predictions suggested that, on average, fewer H2 individuals held at +5 °C (41.7% ± 4.1%) underwent asexual reproduction than anemones held at +2.5 °C (57.3 ± 4.1%) or baseline conditions (47.9% ± 4.1%; Supplementary Table S5). In general, the proportion of H2 individuals undergoing asexual reproduction was overall higher than that for CC7 (H2 baseline mean: 47.9%, CC7 baseline mean: 17.8%). However, CC7 did not experience a temperature-related drop-off in the overall proportion of the population that produced recruits at +5 °C. This difference effectively skewed the size class ratios of both populations (Fig. 2d). Because H2 anemones held at +5 °C experienced a relative decline in the ability to produce new juveniles, but were able to grow to adulthood at elevated temperatures, the average anemone size was larger at +5 °C (12.6 mm^2^ ± 0.46) than at baseline temperature (11.9 mm^2^ ±0.46)(Dunn Test, *p* = 0.043)(Fig. 2d; Supplementary Table 6 and 7). Conversely, CC7 populations held at higher temperatures were primarily comprised of smaller individuals (12.7 mm^2^ ± 0.48 at +2.5°C and 12.2 mm^2^ ± 0.41 at +5°C) in comparison to those held at baseline (14.6 mm^2^ ± 0.58) (Fig. 2c)(Dunn post hoc test, *p* = 0.025, *p* = 0.063; Supplementary Table 6).

**Figure 1.**
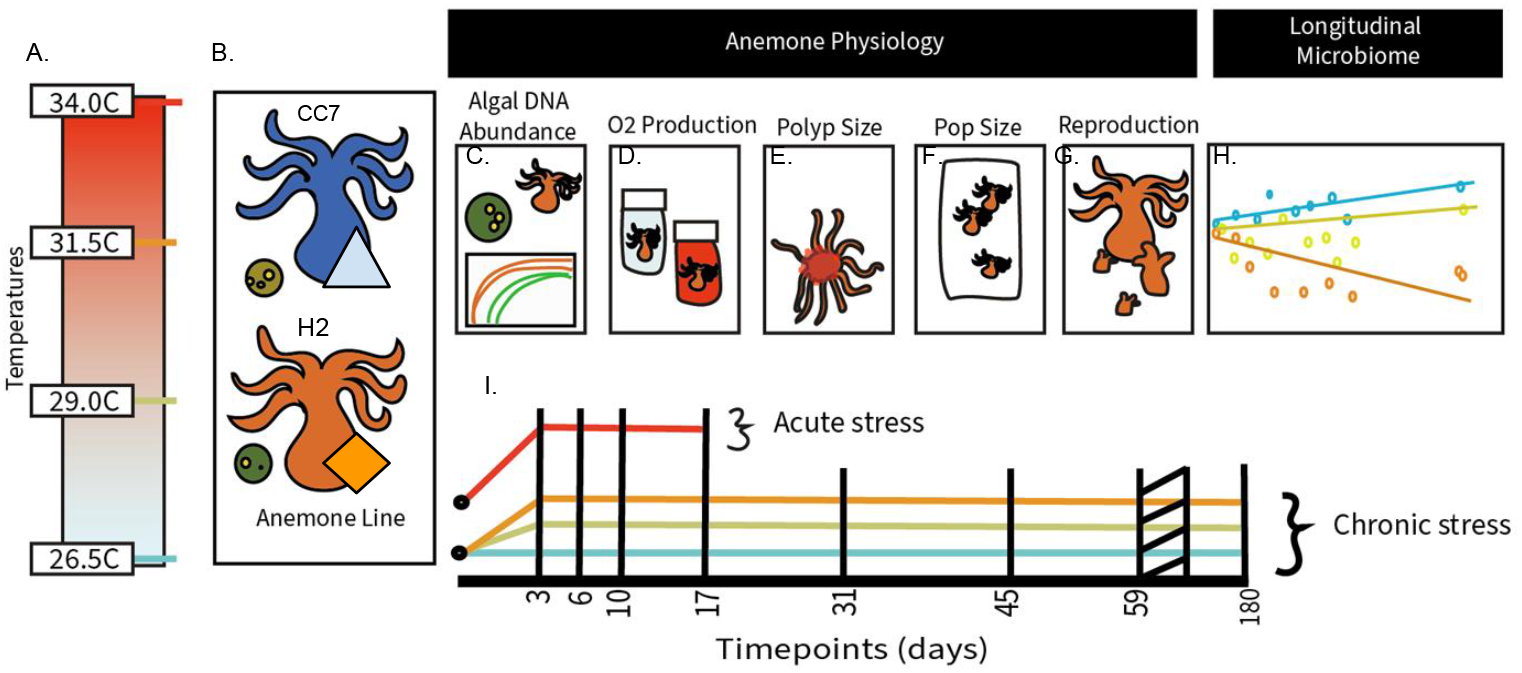
Experimental design. In this study, we test the physiology and microbiome of *Exaiptasia diaphana* across (A.) a heat gradient, from a baseline of 26.5 °C in 2.5 °C increments, to +7.5 °C using (B.) a thermally tolerant anemone line - *Exaiptasia* CC7 with *Symbiodinium microadriaticum* - and a thermally sensitive line - *Exaiptasia* H2 with *Breviolum pseudominutum*. (C.) Anemone photosymbiont to host density was checked 6 times across the course of the experiment, with (D.) showing the photosynthesis to respiration rate, checked at two months. We captured the Population dynamics with (E.) anemone size and (F.) anemone populations three times over the first two months. (G.) Individual anemone reproduction was collected weekly for two months from 144 anemones. (H.) Microbiome data from CC7 and H2 anemones, collected at 0, 3, 6, 10, 17, 31, 45, 59, and 180 days, were used to build linear models for the core microbiome. (I.) The experiment included chronic stress temperatures held for 180 days (baseline, +2.5 °C and +5 °C) and acute stress temperatures (+7.5 °C) held for 14 days after a 3-day heat ramp for all groups.

**Figure 2.**
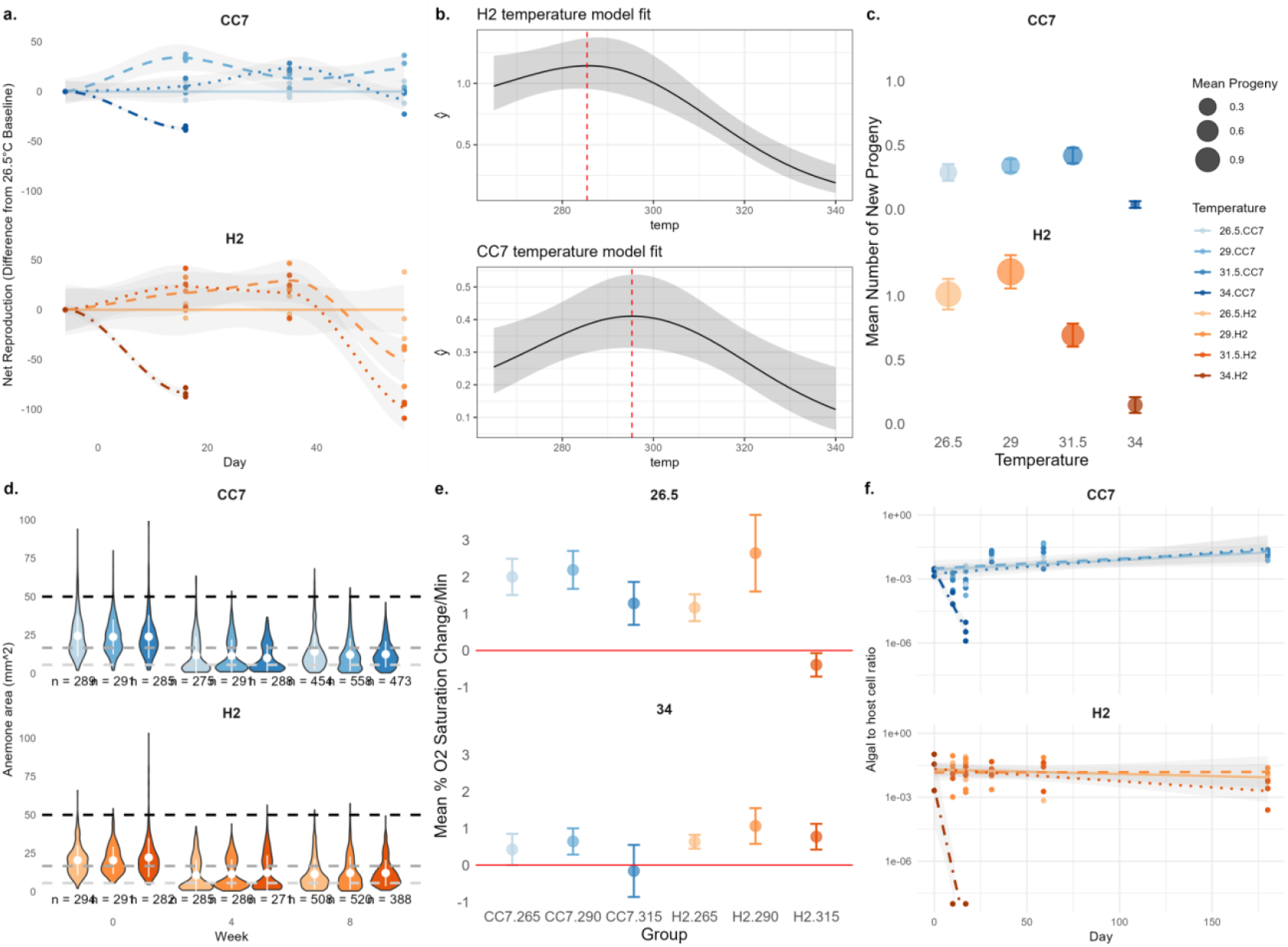
Chronic exposure to elevated stress impacted population and physiology in H2, but not CC7. **a)** Net population reproduction over the first two months relative to the 26.5 °C baseline for CC7 and H2 with loess fits. **b)** GAM predicted fits for temperature effects in reproduction for H2 and CC7 with a vertical line at the predicted optimum for each genet. **c)** Mean pedal lacerates produced per week per anemone for each combination of genet and temperature, with standard errors. **d)** Violin plots showing anemone size distribution, based on area of the oral disk, at timepoints 0, 4, and 8 weeks. Horizontal lines indicate further classification of anemones into small pedal lacerates (below the light gray line), small adults (between the light gray and dark gray lines), adults (between the dark gray and black lines), and large adults (above the black line). We present means and interquartile range using white central dots and lines. **e)** Means and standard error showing photosynthesis to respiration ratios of anemones after 2 months of exposure to baseline (26.5 °C), +2.5 °C, and +5 °C. We conducted the respiration ratio assays at 26.5 °C and 34 °C for both host-algal pairings. **f)** qPCR assays of algal to host ratios across all 180 days with linear model fits.

**Figure 3.**
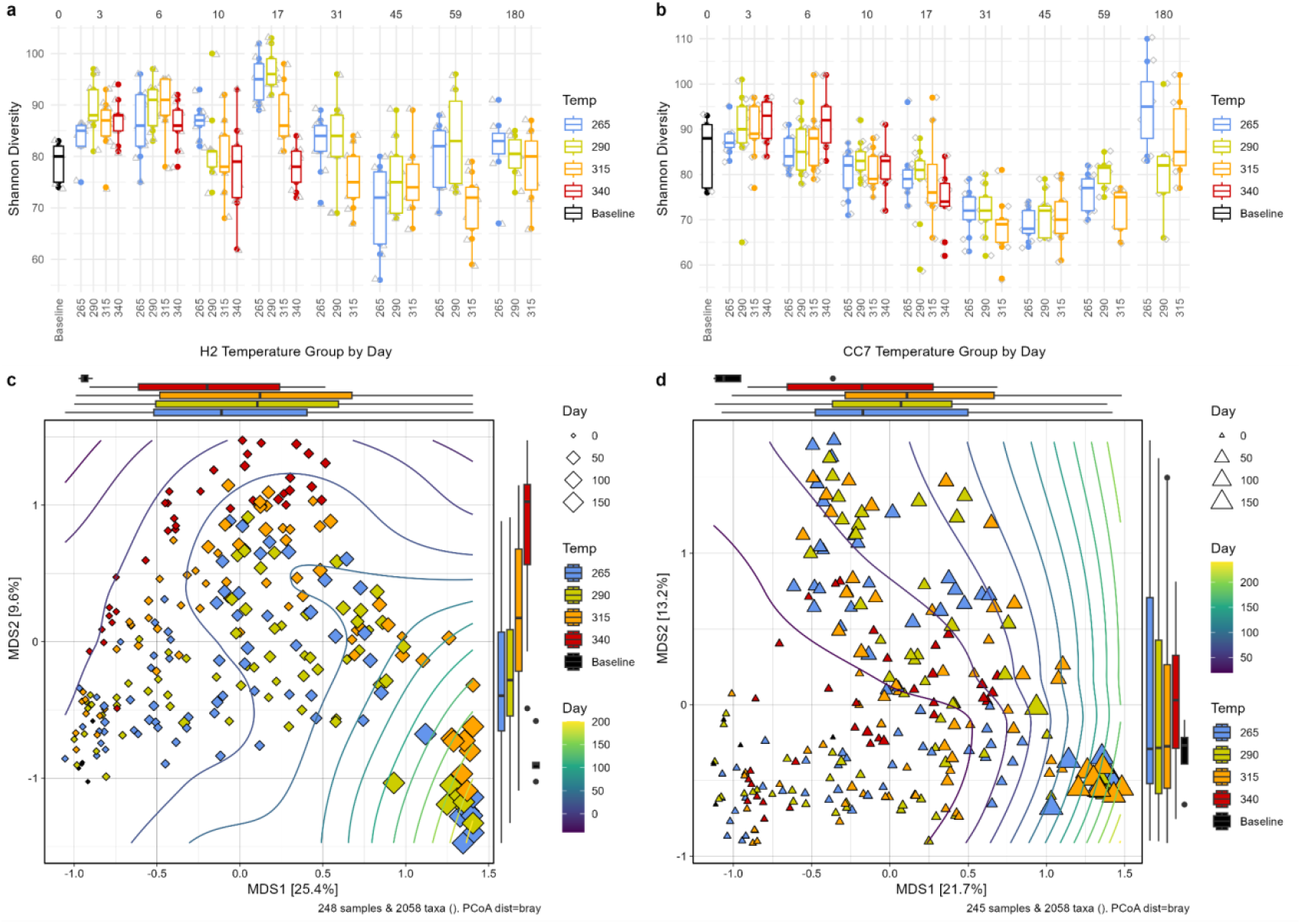
Exposure to elevated temperatures altered the composition but not the richness of the *E. diaphana* microbiome. **a)** H2 Shannon diversity by temperature treatment (color), separated by day (top of x-axis). **b)** CC7 Shannon diversity by temperature and separated by day (top of x-axis). **c/d)** Principal Component Analysis (PCoA) of Bray-Curtis dissimilarity of c) H2 and d) CC7 with points sized by time point and colored by temperature. Contour lines shaded from purple to yellow represent sampling days, from day 0 to 180.

Chronic exposure to elevated temperatures had a negative impact on algal and host physiology in H2, but not in CC7 populations. Photosynthesis to respiration (P:R) ratios measured at baseline temperatures were significantly lower in H2 anemones chronically exposed to +5 °C in comparison to all other groups (ANOVA, *p* = 0.002, Supplementary Table S8). Specifically, net photosynthesis in the H2 baseline, or +2.5 °C, was reduced to 47% of baseline levels when assayed at 34 °C. In contrast, H2 anemones held at +5 °C exhibited the opposite trend, increasing from zero gross photosynthesis at 26.5 °C, with a P:R of -0.39, to 0.79 P:R at 34 °C, which was greater than the mean P:R of 0.52 measured across all other groups. Finally, dark respiration rates for both CC7 and H2 were not impacted by chronic exposure to elevated temperatures, but were affected by assay temperature (Supplemental Tables S9 and S10). Changes in algal performance were not tied to algal densities, as we did not observe significant changes in algal to host ratios (0.017 s/hc ± 0.002, min: 1.7e-4, max: 0.103, Supplementary Tables S11, S12, and S13) across exposure to chronic thermal stress.

Acute exposure to +7.5°C caused the populations to exhibit several negative outcomes in both CC7 and H2. The populations declined, and showed a reduction in algal photo performance in both *E. diaphana* host-algal pairings. Acute exposure also results in CC7 and H2 exhibiting a very low reproduction rate (H2: 11.1% ± 4.3%, CC7: 3.7% ± 2.6%, Supplementary Table S5). Finally, unlike in the +5 °C population, we observed significant and rapid bleaching when we held both host-algal pairings at +7.5 °C (ANOVA, p < 0.001; Supplementary Table S13).

### Exposure to elevated temperatures altered the *Exaiptasia* microbiome

We investigated patterns in the *Exaiptasia* microbiome diversity and community structure from 493 samples collected over 180 days. Our 16S rRNA dataset resulted in an average of 33,858 ± 9,312 reads per sample for a total of 16,692,057 reads. From this read set, we calculated 2,058 Amplicon Sequence Variants (ASVs). The majority of the ASVs belonged to the Alphaproteobacteria (31.8%), Gammaproteobacteria (21.4%), and Bacteroidia (23.2%). Three orders accounted for 44.5% of all sequences, including the Rhodobacterales (18.4%), Chitinophagales (13.6%), and Enterobacterales (12.5%). Critically, we did not observe significant differences in the alpha diversity, measured using the Shannon diversity, of the community between *E. diaphana* pairings or temperature conditions (Fig. 2a-b). Instead, the evenness of the microbial community oscillated over time, where we observed the lowest diversity on day 31 and 45 of the experiment (GAM ANOVA, *p* < 2e-16; Supplementary Fig. 14). Because we see differences in both population-level data and performance between host-algal pairings at chronic and acute exposure, we chose to investigate changes in microbiome composition in each pairing separately. Temperature and time significantly altered the community structure of the *E. diaphana* microbiome in both CC7 and H2 (PERMANOVA *p* < 0.001; Supplementary Table 15). However, the day of sampling or length of exposure, rather than exposure to thermal stress itself, explained the majority of the variation in community structure for both CC7 (48.8%) and H2 (52.7%) host-algal pairings. Conversely, temperature exposure only explained approximately 5.6% of the variation of the community in both CC7 and H2 (Fig. 2a-d). When we investigated the variation in the community structure within sampling day, we observed restructuring of the microbiome related to temperature treatments in both CC7 and H2 populations (pairwise PERMANOVA, *p* < 0.01; Supplementary Table S16). However, exposure to elevated temperatures appeared to have a lesser impact on CC7 than H2 microbial composition. Indeed, pairwise comparisons of the +5°C condition and baseline temperature explained only 2.4% of the variation in the community structure for CC7, but explained 5.9% of the structure variation in H2 (All *r*^2^ < 0.1; Supplementary Table S16). Critically, within-group variance (beta-dispersion) was only significantly different for H2 in the +7.5°C treatment, indicating that the observed differences in community structure were treatment effects and not within-group dispersion (Tukey HSD, *p* < 0.05; Supplementary Table S17).

### Chronic heat significantly modifies the common and core microbiome

Out of 2,058 ASVs, only 46 persistently associated with either H2 or CC7 host-algal pairings. We defined these ASVs as either “core” members of the microbiome (prevalence > 90% within the baseline temperature group) for H2 and CC7 populations or “common” (prevalence > 70% across all samples). We chose these thresholds to capture both the consistent ASVs in the *Exaiptasia* microbiome associated with baseline thermal conditions (i.e., core microbiome) and capture ASVs that are occasionally present at baseline temperatures but reproducibly increase in relative abundance at higher temperatures (i.e., common microbiome). The 46 common and core ASVs accounted for an average of 75% of the relative abundance within each sample (Supplementary Table S18). The vast majority of the common and core ASVs overlapped between host-algal pairings, with 31 out of 46 ASVs overlapping across the H2 and CC7 microbiomes. Of these 31, 24 met the standards for core microbiome in both host-algal pairings, and an additional seven were shared across common thresholds. Eight CC7-specific ASVs were part of the CC7 core or common microbiomes, and seven H2-specific ASVs met these standards.

Of the shared ASVs, 22% (7/31) decreased in both H2 and CC7 under long-term heat stress at +5 °C; three additional ASVs decreased in CC7, and three decreased in H2. In H2, three ASVs from the core and common microbiome increased under long-term heat stress (Supplementary Tables S19-S21). No ASV unique to the CC7 core microbiome increased in response to long-term temperature exposure. In contrast, one ASV within the H2 common microbiome increased in relative abundance at +5 °C, and two members of the H2-specific core microbiome decreased.

All seven ASVs that showed a conserved response between host-algal pairings significantly decreased in relative abundance following chronic exposure to +5 °C (ANOVA, BH adjusted, p < 0.05; Supplementary Tables S20 and S21). These seven ASVs included Nannocystales (ASV16), Pseudomonadales (ASV24), Rhizobiales (ASV28 *Pseudarhensia* sp.), Rickettsiales (ASV73 *Candidatus Megaira* sp.), Enterobacterales (ASV20 *Thalassotalea* sp.), F9P41300-M23 (ASV65), and Francisellales (ASV57 *Francisella* sp.). While ASV16, ASV24, ASV20, and ASV28 all averaged >1% sequences per sample, ASV57, ASV65, and ASV73 averaged under 0.5% of all sequences per sample. In total, these taxa accounted for a cumulative mean of 7.29% of the relative abundance of the community across samples. Therefore, these ASVs were considered low-abundance in both CC7 and H2’s microbiome (Supplementary Table S18).

### Acute and chronic heat stress elicited similar shifts in the microbiome

To compare the changes in the core and common microbiome between chronic and acute exposure to elevated temperatures, we exposed anemones to +7.5 °C (34°C) for two weeks, which resulted in the expulsion of the algal symbionts from CC7 and H2 host-algal pairings (Fig. 2f). In total, acute exposure to +7.5 °C resulted in a significant change in the relative abundance of 17 core and common ASVs in CC7 and 14 core and common ASVs in H2. Nine of these core and common ASVs changed in both CC7 and H2 (linear model, *r*^2^ > 0.35, ANOVA BH adjusted, *p* < 0.05; Supplementary Tables S22 and S23). Declines in +5 °C and +7.5 °C were well conserved. 75% of core and common ASVs that decreased significantly in +7.5 °C exposure decreased in one or both pairings under chronic +5 °C exposure (Fig. 4). This trend was not true for taxa increasing under acute stress; only one ASV increased in both acute and chronic heat stress. Finally, we observed five ASVs belonging to *Candidatus Megaira* (ASV73), *Thalassotalea* (ASV20), *Francisella* (ASV57), *Pseudarhensia* (ASV28), and an ASV within the alphanumeric Gammaproteobacteria order F9P4 1300 (ASV65), which all significantly declined in relative abundance in both pairings and following acute and chronic exposure to elevated temperatures over time.

**Figure 4.**
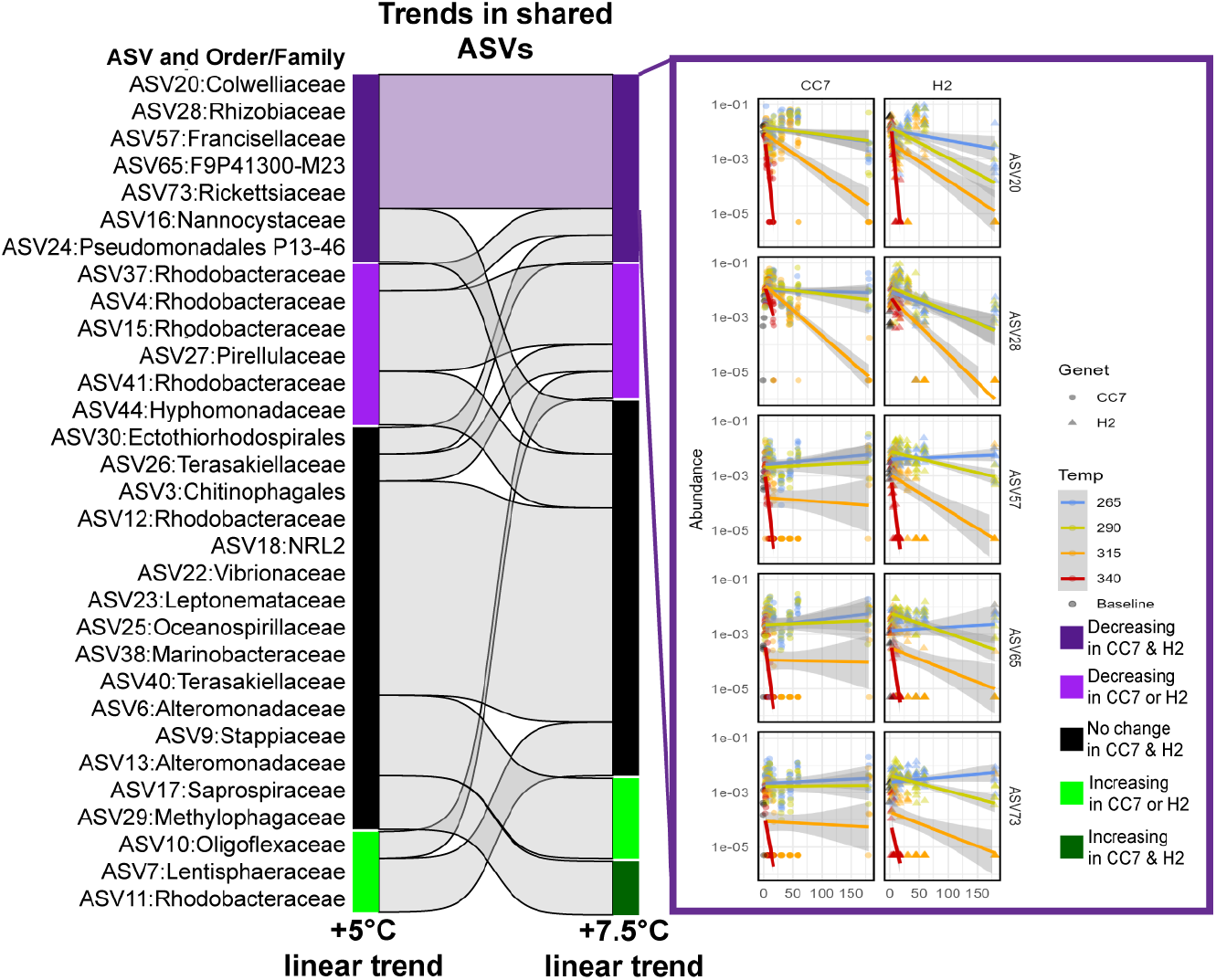
Five core and common taxa shared a conserved response to acute and chronic exposure to stress. Sankey plot of trend conservation across chronic and acute stress. All shared ASVs and their associated trend in long-term (chronic) +5 °C exposure (left) and short-term (acute) +7.5 °C exposure (right). Categories represent an increase in both (dark green), one (green), and a decrease in both (dark purple) or one (purple). Non-significant trend groups are in black. Linear trends for the five conserved downward-trending ASVs are present on the right-hand side across 180 days.

To identify indicator taxa that could be driving dysbiosis of the microbiome, we tested all ASVs with conserved chronic and acute responses for significant differences in log-relative abundance at +5 °C after seven and fourteen days. These time points occur before any population-level decreases were apparent in H2. Additionally, for the acute groups, we mapped microbial relative abundance changes to the day on which we completed the heat ramp, and to three days post-heat ramp. These timepoints correspond to the timeframe during which the transcriptomic response occurs in *E. diaphana*. In the temperature-sensitive H2 host-algal pairing, the relative abundance of ASV73 *Candidatus Megaira* was distinct from the baseline temperature group by one week at +5 °C. By two weeks, ASV73, ASV57 *Francisella*, and ASV65 F9P41300 were also significantly lower than in base temperatures in both CC7 and H2. ASV20 *Thalassotalea* was significantly different in H2 at one week, and in CC7 at two weeks, but was not significant at two weeks in H2 (Kruskal-Wallis & Dunn post hoc tests, p < 0.05, Supplementary Tables S24 and S25, Log_2_ fold change on S26).

Under acute stress, ASV20, ASV57, and ASV73 were all significantly lower in both H2 and CC7 by three days, but not before three days. ASV65 was significantly lower in CC7 by three days; however, relative abundance changes were below the threshold of significance for H2 (KW and Dunn post-hoc tests, p < 0.05, Supplementary Tables S27 and S28). In H2 only, ASV28 *Pseudarhensia* was lower by three days in acute stress.

Based on these results, we categorized ASV20 Thalassotalea, ASV73 *Candidatus Megaira*, ASV57 *Francisella*, and ASV65 F9P41300 as early heat stress indicator taxa.

### Exposure to elevated temperatures altered microbiome networks more strongly in H2 than CC7

To identify changes in microbial co-occurrence patterns, we constructed networks for each host-algal pairing and each temperature combination using all taxa with more than 1000 reads (177 taxa). Of these, between 114 and 125 ASVs had adequate representation across samples within a given temperature for network inclusion. For the thermally tolerant CC7, network restructuring was limited, with microbial networks remaining well-correlated across all subacute temperatures (Spearman’s ρ: 0.65-0.77; Fig. 5, and Supplementary Table S29). In contrast, the thermally-sensitive H2 showed greater network dissimilarity between the baseline and +5 °C treatments (Spearman’s ρ = 0.56). In CC7, abundances were better correlated between +2.5 °C and baseline; H2 microbial networks were as divergent between baseline and +2.5 °C as between +2.5 °C and +5 °C (Fisher’s Z, p = 0.37; Supplementary Table S30). This difference suggests that divergence occurs at lower temperatures in the H2 microbiome. These changes resulted from higher within-network correlation; edges increased by 14% in CC7 at +5 °C relative to baseline, 27% in H2 (see Supplementary Table S31). Furthermore, the number of connections per ASV increased significantly in the H2 +5°C network compared to the baseline, a trend that was not significant in CC7 (Supplementary Table S31). This contrast indicates a more pronounced restructuring of microbial interactions in the thermally sensitive pairing.

**Figure 5.**
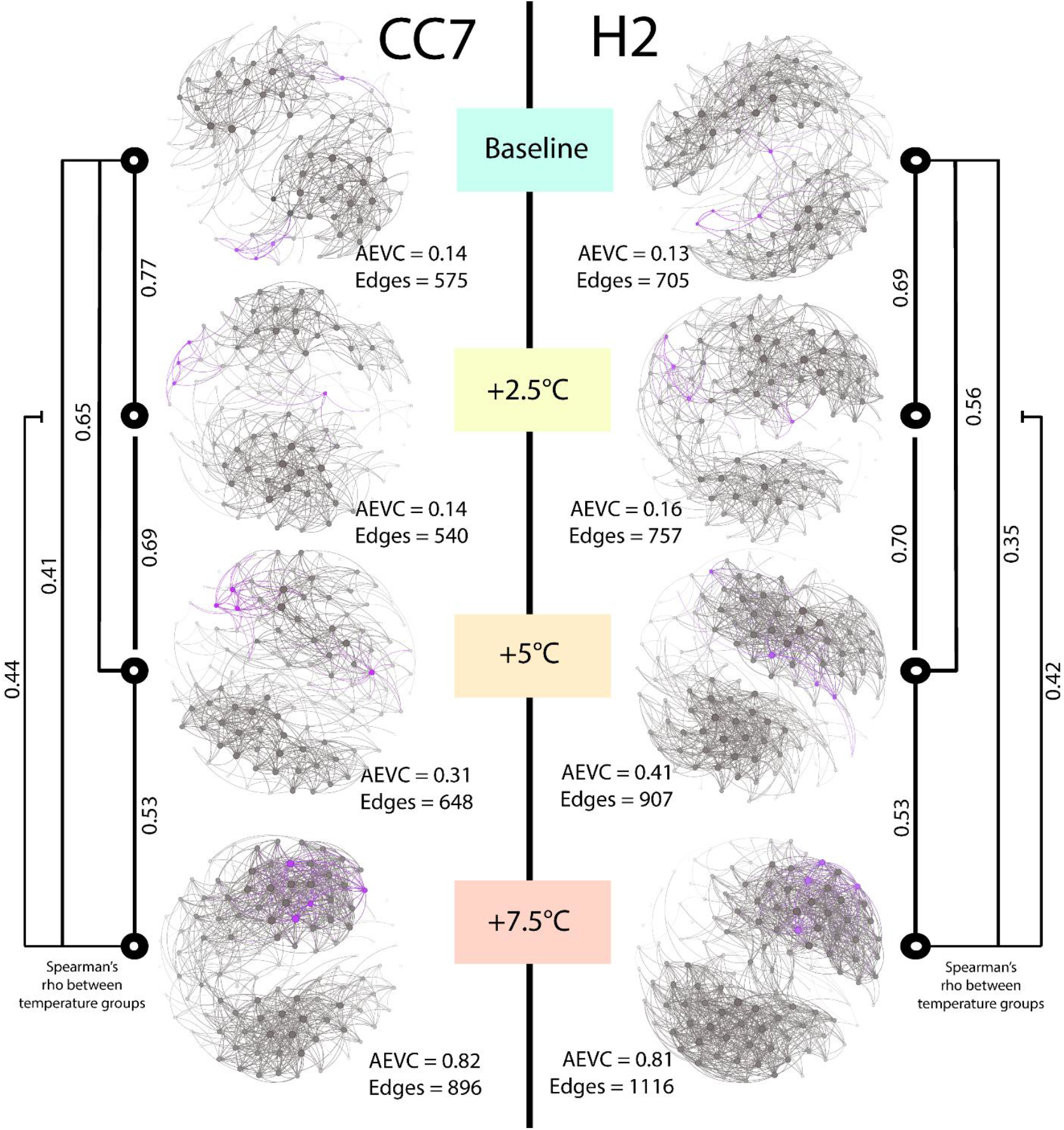
Heat indicator taxa grow more connected at increasing temperatures. Network maps of ASVs sized by number of connections (*r*^2^ > 0.4) in each genet and temperature combination (edge darkness based on association strength). For each network, the average eigenvector centrality (AEVC) and the number of connections (Edges) are given. Additionally, we present Spearman’s rank correlation (*ρ*) between all ASV associations (n = 15,753) for each combination of temperatures within each host. We display conserved early heat indicator ASVs in purple (ASV20, ASV73, ASV57, and ASV65).

**Figure 6.**
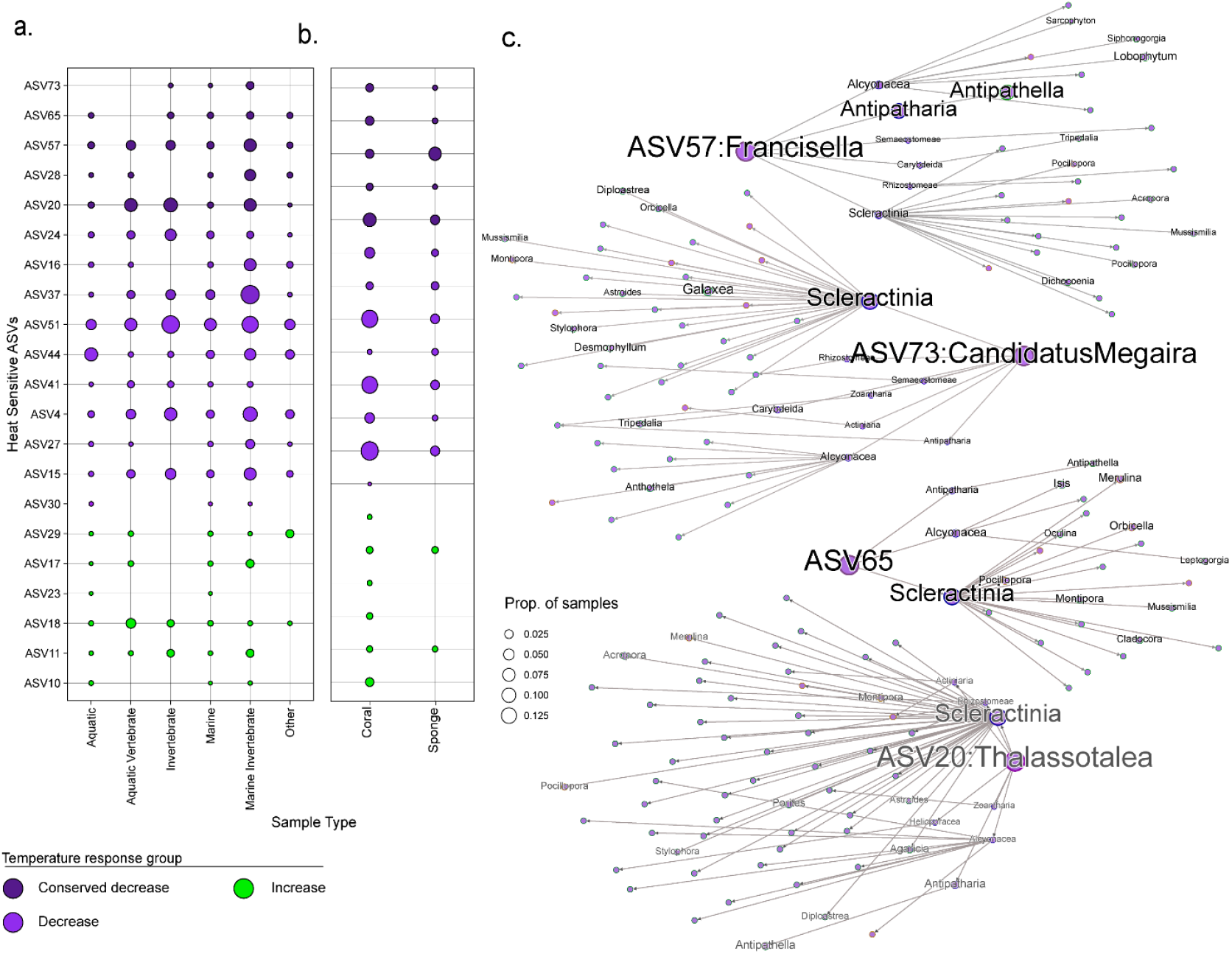
Heat-sensitive ASVs are present in other cnidarians. a) Prevalence (percent of samples) of an ASV within a given sample type. Point size indicates the relative abundance, and point color represents the heat group (conserved decreasing, decreasing, or increasing) from this study. b) Panel two includes only annotated coral and sponge metagenome samples for comparison. Within panel one, these are included within marine invertebrates. c) Networks of genus presence across cnidarian orders and genera (names for the top ten genera in terms of ASV prevalence displayed) for the genera of the top four indicator taxa, *Thalassotalea, Candidatus Megaira, Francisella*, and genus-level divergence from ASV65 F9P41300-M23.

While in both CC7 and H2, +7.5 °C was the network with the lowest correlations compared to all the other temperatures, the correlation between +5 °C and +7.5 °C in H2 is of a similar magnitude as between baseline and +5 °C (Spearman’s ρ = 0.53 vs. 0.56). This disparity is not observed in the CC7 microbiome, where the conservation between baseline and +5 °C is substantially higher than between +5 °C and +7.5 °C (Spearman’s ρ = 0.65 vs. 0.53). Of all comparisons between host-algal pairings, chronic stress produced the most similar networks (Spearman’s ρ = 0.56 vs. 0.48-0.49)

Overall, changes in modules for H2 and CC7 were not strongly associated with temperature. Within the modules of H2 and CC7, we characterized the Jaccard distance between all modules across all six networks. We identified the highest Jaccard values (i.e., the fewest number of changes to the module) for all modules within the networks with increased temperature. High maximum Jaccard similarity between temperature groups indicates that the membership of cooccurrence modules is stable across temperatures. Comparing the H2 baseline to H2 +2.5 °C and the CC7 baseline showed higher group continuity than H2 at both chronic and acute stress (Dunn test, p-value < 0.05; Supplementary Tables S33 and S34). This similarity suggests that highly correlated taxa under baseline temperatures in H2 remain highly correlated in H2 +2.5 °C, but to a lesser extent in H2 +5 °C.

We tracked the heat indicator taxa across all modules to identify patterns of association. If temperature-sensitive ASVs were found across differing modules at baseline temperatures, but converged in higher temperatures, this would suggest that temperature sensitivity was their main shared trait. If temperature ASVs respond in tandem at stable temperatures, these may represent a true functional unit. We observe ASV57 *Francisella*, ASV73 *Candidatus Megaira*, and between one and three ASVs from the group F9P41300-M23 (ASV65, ASV89, ASV105) consistently in a stable module at temperatures below +7.5 °C (See Supplementary Table S32). The F9P41300-M23 clade was the only one in this module where the specific ASV was flexible. In CC7 +5 °C, this module included ASV62 *Hyphomonas*, but no other deviations were present. Under acute temperature stress, this module breaks across multiple modules in both H2 and CC7. ASV20 *Thalassotalea* varied separately from this module, indicating a separate functional grouping.

At baseline temperatures, these ASVs (ASV57, ASV65, ASV73) operate primarily within an independent module, which is poorly connected to the rest of the network, with average eigenvalue centrality of 0.119 CC7 and 0.093 in H2. At +5 °C, eigenvector centrality for these nodes increases dramatically (eigenvector centrality values = 0.302 in CC7 and 0.454 in H2; Supplementary Table S31). In +7.5 °C, they are among the most connected nodes in the network (eigenvector centrality values = 0.755 in CC7 and 0.766 in H2). Since we construct these networks using only positive edges with a high *r*^2^ (greater than 0.4), greater connectedness at higher temperatures indicates that other connected taxa within these networks are likely decreasing, particularly at levels below the significance threshold in CC7 and H2.

### Indicator taxa identified in *Exaiptasia* are widespread in other cnidarians

To assess the broader relevance of our findings, we queried our core ASVs against global databases. We detected core members of both the H2 and CC7 microbiome in datasets on Integrated Microbial Next Generation Sequencing (IMNGS) and in a global 16S rRNA survey of coral microbiomes^1^. Heat-sensitive ASVs varied in prevalence across microbiomes, with some absent from IMNGS and therefore specific to our study. Additionally, we found other ASVs in up to 20% of all annotated coral IMNGS 16S rRNA samples (range: 0.04% to 12.95% prevalence per ASV; Supplementary Table S35). Overall, marine invertebrates contained higher average proportions of our ASVs than other sample types, and coral metagenomes had significantly higher prevalence of taxa that decreased with heat than increased (KW and Dunn post-hoc tests, p < 0.05; Supplementary Tables S36, S37, and S38). Heat indicator taxa (ASV20, ASV73, ASV57, and ASV65) ranged in frequency from 1.25% of coral samples to 5.76% (ASV20) (Supplementary Table S39). As we can identify these heat-impacted ASVs across a meaningful proportion of IMNGS coral samples, we then identified specific cnidarian genera in which they were found. When we re-analyzed a global cnidarian microbiome dataset, we found that the heat-impacted ASVs identified in our study were present in an average of 9 other cnidarian species (Supplementary Table S41). These findings suggest that the specific heat-sensitive taxa identified in *Exaiptasia* are not unique to this model system but are present across a wide range of cnidarian hosts, highlighting their potential as bioindicators of thermal stress.

## DISCUSSION

### The core and common microbiome is a small part of the Exaiptasia microbiome, with some of the core having low relative abundances in each sample

Cnidarian microbiomes are an increasingly well-sequenced ecological niche; however, our understanding of temperature association is limited by the complexity of the system^1,24^. In addition to a high starting diversity, variation in photosymbiont community and host thermal tolerance has profound impacts on microbial communities before, during, and after a stress event^31,33,35,36^. Within this study, we find that the core microbiome of *Exaiptasia* host-algal pairings, as defined by relaxed prevalence standards -90% in baseline samples or 70% across all samples for one pairing, made up an extremely low proportion of the ASVs within the anemone tissues. In some of these core members, the mean relative proportion of sequences was relatively low (0.2% for ASV73 *Candidatus Megaira*, 0.6% for ASV40 *Rhodovulum*, and 0.4% for ASV57 *Francisella*). These trends may be substantially more pronounced in cnidarians with higher surface area to tissue ratios, such as corals and hydrocorals^37^. Taxa that are in fact core microbiome associates may run the risk of being stripped from analyses for low abundances or labeled as rare, especially in studies rarefying to low thresholds.

### Heat impacted the core microbiome regardless of *Exaiptasia*’s physiological stress

The CC7 *Exaiptasia* did not experience physiological declines at +5 °C, but did experience contemporaneous shifts in some core microbiome members with the H2. This disparity suggests that decreases in core microbiome members can be tolerated, but may be indicative of encroachment on thermal limits. This difference is also highlighted by the apparent fitness of H2 at +2.5 °C, which displays some core microbiota reductions over a longer time horizon than +5 °C despite physiological stability. As CC7 and H2 experience different stress thresholds, this conserved group response is of significance.

We see substantial diversity in response within orders in the *Exaiptasia* core microbiome. Chitinophagales, Rhizobiales, and Enterobacterales all include ASVs that respond differently to chronic and acute heat exposure. It is now well understood that 100% V4 identity is a reasonable threshold to avoid lumping species, but continues to underrepresent the diversity of a sample. With our large sample size, we resolve a reasonably compact core microbiome that includes ASVs across many orders and families.

### Changes in the relative abundance of a five ASVs is conserved under both acute and chronic stress in CC7 and H2 strains

Within the four heat indicator ASVs impacted in both host-algal pairings and under acute and chronic stress, each represents a unique bacterial genus of interest. *Thalassotalea*, an unknown ASV65 from F9P41300-M23 (closest NCBI Blast sequence match 89% *Pseudomonas fluvialis* NR_159318.1), *Francisella*, and a Rickettsiae *Candidatus Megaira* (closest NCBI Blast sequence match 94% *Rickettsia tillamookensis* NR_181615.1). All of these ASVs are identical to ASVs from an earlier paper on *Exaiptasia* thermal tolerance.

The first of these genera, *Thalassotalea*, is a genus of aerobic complex carbon degraders with isolates cultured from deep water corals, shallow water corals, and oysters^38^. While some species within the genus are chitin degraders, this is not universal^39^. Researchers have not yet described the location of this genus within host tissues, but it may be a member of the gastrovascular community.

The genus *Francisella* is best known for its pathogenicity in humans, but has also evolved as a nutrient-provisioning symbiont within ticks on multiple occasions^40^. In the marine environment, it has been found as a ciliate endosymbiont^41^ and as a fish pathogen^42^. Within photosymbiotic corals, *Francisella* are regularly observed, although their position as an endosymbiont or opportunistic pathogen is unclear^43^.

The *Rickettsia* group is one of ongoing interest in marine intracellular symbioses and parasitology. While regularly found in coral samples and other cnidarians, there is uncertainty about whether they may be facultative endosymbionts or parasites weakening hosts in the face of different pathogens. For algae, *Candidatus Megaira* is a potential defensive symbiont^44^. The widespread *Aquarickettsia* is a marine invertebrate parasite that may persist in nutrient-dense environments^45^. *Megaira* and *Rickettsia* lineages may represent exclusively pathogenic genera or, like many marine species, may include a mix of strategies^46,47^.

Given the high similarity in trajectories of abundances between the *Candidatus Megaira, Francisella*, and ASV65 Gammaproteobacteria lineage, these may represent a group of intercellular parasites or endosymbionts lost in temperature stress conditions. Their presence may be because a) they persist primarily on excess photosynthate, b) they are each independently incapable of surviving the host environment at higher temperatures, or c) they represent nutritional symbionts no longer worth provisioning in stressed hosts.

## CONCLUSIONS

The microbiomes associated with invertebrate hosts risk destabilization in acute and chronic heat stress. In *Exaiptasia*, acute and chronic stress impact different members of the core and common host-associated microbiota, with a sensitive host genotype displaying more significant network alterations than a tolerant host. Despite these distinctions, host health was not an arbiter of network destabilization, as the tolerant host experienced shifts at elevated temperatures as well. We additionally demonstrate that these same ASVs are common across sample types within broader marine databases, including both conphyletics and non-Cnidarians. We also highlight that the strongest temperature signal may come from potential tissue symbionts of *Exaiptasia* CC7 and H2.

## METHODS

### Experimental design

1444 Exaiptasia from CC7 and H2 host-algal pairings were raised under 12:12 light:dark cycle (PAR 90+/-20) at 26.5 °C in 35 ppt artificial saltwater (Red Sea). At the start of the experiment, anemones were split into 26 replicate populations of 50 anemones each (13 H2 and 13 CC7 populations held in 4L tanks). An additional 24 six-well plates of H2 and CC7 (12 plates per host-algal pairing) were prepared with one anemone per well. All tanks and plates were held at baseline for three days, then split into three tanks per genet/temperature combination (26.5°C, 29°C, 31.5°C, 34°C). One tank of CC7 and H2 was reserved for producing baseline data. Temperatures for each population were ramped across five 12 hr increments to reach their final holding temperatures (0 °C/12 hr for baseline, 0.5 °C/12 hr for +2.5 °C, 1 °C/12 hr for +5 °C, 1.5 °C/12 hr for +7.5 °C). Throughout the experiment, partial water changes of 1 L occurred daily. Full water changes (3L) occurred three times per week, 6 hours after feeding. Artificial seawater was prepared using Red Sea artificial sea salt, sterilized using a UV sterilizer, and maintained at a constant temperature of 30-32 °C. Water changes for six well plates occurred daily. Tank changes occurred 3x over the first two months of the experiment (19-21 days apart) with tanks UV sterilized before anemone addition. After 2 months, as sampling was less frequent, feeding and water change frequency were reduced to 2x/wk.

### Imaging

To quantify shifts in individual anemone size, anemone populations were imaged three times per week using a light table with a document camera (JOYUSING V500). Photographs from three time points (start, one month, and 2 months) were hand masked for segmentation of each anemone. Exaiptasia within these masks was isolated using Python’s SKImage (measure.label). The top-down anemone oral disc area was calculated for each labelled mask using the pixel count to give the size distribution.

### Population and reproduction hand counts

Hand population counts were measured at the time of tank changes (3x) across the first two months of the experiment. Briefly, each anemone was gently dislodged from the tank and counted. To avoid tank overpopulation, only 50 adult anemones were moved into the new tank, and the remainder were discarded, as net population was calculated three weeks after each move; net reproduction rates are directly comparable. To estimate asexual reproduction rates, we counted the number of pedal lacerates per individual in anemones held in six well plates.

### Respiration Assay

At two months, single anemones held in six well plates were moved into filtered autoclaved artificial salt water at 35 ppt in 10 mL vials with oxygen-reactive PreSens labels. Anemones, along with one oxygenated and one deoxygenated control (prepared according to PreSens instructions), were placed onto a PreSens reader mounted on a rotating table in 26.5 °C or 34 °C in the dark. After reader stabilization (∼15min), oxygen consumption was monitored for 60 min in the dark. After dark measurements were collected, lights (80 PAR) were turned on, and after an additional stabilization period (∼1 hr), oxygen measurements were collected for net photosynthesis. Anemones were flash frozen in liquid nitrogen and stored at - 80 °C until we quantified total protein using a BSA kit.

### Host-Symbiodinium qPCR assay

To quantify algal density within the host tissue, we assessed changes to the ratio of algal DNA to host DNA using qPCR. A genus-specific ITS2 primer set was used to quantify algal copy number^48^, and host DNA was quantified using translation elongation factor 1 or Ef1-a^49^. qPCRs were performed on a QuantStudio3 in triplicate 20 µl reactions using 10 µl of Powertrack Sybr green master mix, 1 µl of template DNA (see DNA extraction below), and 1 µl of 10 µM Forward and Reverse primers. Both algal primer sets used the same thermal conditions: initial denaturation for 2:30m at 95 ºC, followed by 40 cycles of 95 ºC for 0:15 s, 62 ºC for 0:30 s, and 72 ºC for 0:30s. For host DNA, thermal conditions were as follows: initial denaturation for 2:00 m at 94 ºC, followed by 40 cycles of 94 ºC for 0:15 s, 60 ºC for 0:30 s, and 72 ºC for 0:30 s. A standard 10-fold dilution series from 10^−1^ to 10^−7^ ng/µl of purified gene of interest was run in duplicate on all plates and used to calculate the copy number.

### Statistical Analysis

The effects of temperature, genet, and time on anemone net reproduction in tanks were analyzed using Generalized Linear Models (GLMs) with a Gaussian distribution on log-transformed reproduction counts (implemented in the “mgcv” package and “gratia” package)^50^. Analysis of Variance tests (ANOVA) were used to assess the relative contributions of temperature factors to the model. We employed Generalized Additive Models (GAMs) with a negative binomial distribution to model the reproductive rate of individual anemones, fitting smooth terms for time (day) and temperature. The GAM ANOVA (“mgcv”) was used to assess the relative contributions of temperature to the model. Anemone size (area) was compared across conditions using a Kruskal test and Dunn post hoc comparison with Benjamini-Hochberg adjustment for multiple comparisons.

The ratio of symbiont-to-host cells, determined by qPCR, was log-transformed to meet model assumptions. A GAM was used to assess the effects of collection date, temperature, and genet, including their interactions, on the host-symbiont ratio with a GAM ANOVA post-hoc test to assess the significance of factors. P:R ratios were computed as gross photosynthesis (light oxygen change (% saturation) - dark oxygen change (% saturation)) over respiration, then the factor significance of temperature group, genet, and measurement temperature were checked with analysis of variance (ANOVA on GLM results). The same features were applied to a model of protein-adjusted respiration.

### Microbiome sampling

Replicate anemones (n=3-5) were sampled from each of the six populations per temperature treatment at days 3, 6, 10, 17, 31, 45, 59, and 180 for DNA extraction. To capture microbial taxa associated with the ASW, we filtered 1 L of ASW through a 15 µm filter in triplicate at each time point. All samples were preserved in 500 µL of Zymo DNA/RNA shield, snapfrozen in liquid nitrogen, and then stored at -80 °C until DNA extraction. DNA was extracted from all samples. DNA was extracted from whole anemones and water filters using a Zymobiomics DNA Miniprep kit (D4300) with the following modification: beating was conducted using an MP Biosystems FAST Prep. It consisted of 5 cycles of 60 seconds at 6 m/s, with a 5-minute cooling period on ice between cycles.

### *16S rRNA* Library preparation and sequencing

The V4 region of the 16S rRNA gene was amplified following the Earth Microbiome Project protocol with the following modifications: the PCR cocktail consisted of 12 µL of Invitrogen Hot Start Platinum Taq, 9 µL of water, 0.375 µL each of the forward and reverse primers using the barcoded primer set 515FY, 806RB from the Earth Microbiome Project (Parada 2016, Apprill 2015)^51,52^, and 0.8 µL of DNA template. PCR conditions were: 94°C for 3 min followed by 38 cycles of 94 °C for 45 s, 50 °C for 60 s, and 72 °C for 90 s, and a final extension period of 72 °C for 10 min. PCR reactions were run in triplicate per sample. Triplicate PCR products were pooled by sample and then pooled to equal molar concentration per sequencing library. In total, we generated three sequencing libraries, which were purified using gel extraction (Monarch DNA). Sequencing was performed at the University of California, Davis Genome Center using an Illumina MiSeq to produce 2 × 250 base pair (bp) reads.

### Processing of the 16S rRNA amplicon sequencing data

Resulting 16S rRNA samplicon data were processed using the DADA2 v 1.34 pipeline in R (v 4.4) to determine amplicon sequence variants (ASVs)^53,54^. Briefly, sequences were filtered and trimmed (“filterAndTrim(truncLen=c(210,160), maxN=0, maxEE=c(2,2), truncQ=2, rm.phix=TRUE”), error corrected (“learnErrors”, “dada”), and merged (“mergePairs”). Only reads between 230 and 270 bp were retained. Chimeric reads were removed (“removeBimeraDenovo”), then taxonomy was assigned using the Silva v 138.1 database modified to include ten additional sequences from known isolates(“assignTaxonomy”)^55^. 27132939 sequences were present after quality filtering and taxonomic assignment. Unclassified ASVs and ASVs assigned to mitochondria and cyanobacteria were removed using the subset taxa function in phyloseq (v1.42), after which 17568069 sequences remained^56^. We used the decontam package (v1.18) to remove contaminants in two rounds: first, we removed ASVs based on differential abundances in PCR and extraction controls (17544042 sequences remaining), and second, based on the differential abundance of ASVs in ASW samples (17136484 sequences remaining)^57^. Finally, samples with fewer than 10,000 reads were removed from the dataset, resulting in 493 samples (16692057 sequences remaining). To verify that this resulted in adequate sequence coverage, rarefaction plots (“rarecurve()”) were generated in vegan v2.6.6^58^.

#### Microbiome diversity

To quantify shifts in the *Exaiptasia* microbiome as a function of thermal treatment, sampling time point, and *Exaiptasia* host-algal pairing, we used the “phyloseq” package to estimate species richness and compare taxonomic profiles across samples. First, we determined Shannon diversity and quantified the number of observed ASVs using the “estimate_richness” function. Shannon diversity was then modeled with a GAM of smoothed time, temperature, and genet, with a GAM ANOVA applied as a post hoc test to identify the significance of fit features.

The raw ASV counts were normalized to relative abundance and used to calculate a Bray-Curtis distance matrix, which informed a separate PCoA analysis for CC7 and H2, displayed with microViz^59^. We used the “betadisper” function to calculate beta dispersion within temperature treatments separately for each *Exaiptasia* host-algal pairing and compared the resulting values using a Tukey’s HSD test. We calculated a PERMANOVA using the “adonis2” function in the “vegan” package to compare multivariate sample distances across treatments, sampling time points, and *E. diaphana* pairing. Pairwise differences were determined using the pairwise.adonis2 function in the pairwiseAdonis package (v 0.4.1)^60^.

### Core microbiome

To determine the core microbiome and common microbiome, we utilized the “microbiome” package^61^. Because the two *Exaiptasia* host algal pairings are known to have distinct microbiomes^62^, we defined the core and common microbiome separately for each host-algal pairing. The core microbiome included ASVs with a presence in more than 90% of all samples held at 26.5 °C across the entire experiment. The common microbiome included ASVs present in at least 70% of all samples at a relative abundance of at least 0.1% across 26.5 °C, +2.5 °C, and +5 °C. The early common microbiome included ASVs present in at least 70% of all samples from 0 days to 17 days for each *E. diaphana* lineage. We chose to include the common and early common microbiome to ensure we could detect taxa that were rare under normal conditions but increased in higher temperatures.

### Linear model testing

Core or common ASVs for both threshold H2 or CC7 were used for linear model testing. Specifically, each ASV was log-transformed, then a linear model was generated (“lm()”) based on the interaction of temperature and collection date in non-acute temperatures (baseline, +2.5 °C, +5 °C). Plots were checked with fit diagnostic plots in base R(“plot()”), and analysis of variance was computed for each model (“anova()”), and p-values were adjusted using a Benjamini-Hochberg adjustment to reflect the number of comparisons (“p.adjust(method = “BH”)”). A p-value of < 0.05 in either the temperature factor or the interaction factor, and an *r*^2^ value of > 0.15 were the thresholds for inclusion. The directionality of the relationship was recorded for cross-comparison.

All ASVs that met a threshold of 70% prevalence in samples from the first two weeks at temperature for each host-algal pairing (“core_members(Genet, detection = 0, prevalence = 70/100)”) were used for +7.5 °C linear models. Selection criteria were the same as those for long-term linear models, but with a more stringent explanatory threshold (*r*^2^ > 0.35), as there was less time for long-term fluctuations.

### Network analysis

Networks were computed in Sparse Cooccurrence Network Investigation for Compositional (SCNIC) for modules and networks (“within” and “modules”) using the top 214 ASVs within the dataset^63^. Modules were removed from the analysis if r < 0.4. Spearman’s *ρ* was computed for connection similarity across all ASV-ASV combinations within each genet temperature combination, and Fisher’s Z was calculated for each Spearman’s *ρ* combination for each genet. The number of connections of each node was compared with a Kruskal-Wallis test and Dunn pairwise comparisons. Module similarity was determined by calculating a Jaccard matrix of all modules across temperature and host-algal pairings. The maximum similarity by module was computed for each group relative to the CC7 baseline and the H2 baseline. The average of these maximum Jaccard distance scores for each module was compared with a Kruskal-Wallis test and Dunn pairwise comparisons. All other network statistics were computed in Gephi.

### Querying IMNGS & McCauley datasets

To determine if the members of CC7 and H2’s core and common microbiomes are present in other datasets, we uploaded the ASVs from each of these taxa to the IMNGS web server. We used the ‘paralleler’ function with all available database accessions to search for closely related sequences (minimum length of 200 bp and minimum identity 99%). Datasets were retained in our analysis if they included at least one ASV that had a minimum relative abundance of 1% in the dataset. Data was reduced to sample types with a minimum of 1% incidence, then combined by group (marine, aquatic, invertebrate, vertebrate, unknown). Mean abundances were compared between heat-impacted taxa and across sample types using Kruskal-Wallis and Dunn post hoc test (“kruskal.test()” and “dunn_test()”).

The global cnidarian V4 phyloseq object from McCauley et al. 2023 was downloaded from the supplementary materials of the paper. The reference sequences from the object were pulled and retrained against a Silva NR138 database with the core microbial ASVs from core microbiome delimitation included. After retraining, samples with fewer than 1000 sequences per sample were removed from the dataset. The resulting count table was normalized to generate a relative abundance table and subset to “species” level identities matching ASVs from the *Exaiptasia* core microbiome. Relative abundance of heat-impacted and unaffected ASVs was compared with a Kruskal-Wallis test, and networks of cnidarian taxa with these ASVs were constructed with ggnetwork.

## STATEMENTS AND DECLARATIONS

### Data Availability

All data generated and codes used in analysis are available upon request.

### Contributions

KMM, SW and EMS were responsible for initial study design. KMM was responsible for microbiome data collection, data analysis and manuscript preparation. SW was responsible for image processing, segmentation and analysis, and assisted in the analysis of physiological metric data and manuscript refinement. SM was responsible for qPCR assay design and finalization. KMM, HHW, SaAM, SuAM, SM, and MJWS were responsible for maintenance of the experiment, sample collection, physiological testing and manuscript refinement. EMS was responsible for project oversight, funding, manuscript editing and approval of the final manuscript. All authors read and approved the final manuscript.

## ACKNOWLEDGEMENTS

This work was supported by a National Science Foundation under Grant No. DBI-2214038. The work was carried out as part of University of California, Merced’s BII-INSITE.

## REFERENCES

1. McCauley, M., Goulet, T. L., Jackson, C. R. & Loesgen, S. Systematic review of cnidarian microbiomes reveals insights into the structure, specificity, and fidelity of marine associations. Nat. Commun. 14, (2023).

2. Bent, S. M., Miller, C. A., Sharp, K. H., Hansel, C. M. & Apprill, A. Differential Patterns of Microbiota Recovery in Symbiotic and Aposymbiotic Corals following Antibiotic Disturbance. mSystems 6, 1–18 (2021).

3. Perez, S. F., Cook, C. B. & Brooks, W. R. The role of symbiotic dinoflagellates in the temperature-induced bleaching response of the subtropical sea anemone Aiptasia pallida. J. Exp. Mar. Bio. Ecol. 256, 1–14 (2001).

4. YKL, JHL, HKL. Microbial Symbiosis in Marine Sponges. J. Microbiol, 39(4):254–264 (2001).

5. Arai, M. A Functional Biology of Scyphozoa. Chapman & Hall, (1997).

6. Kruse E., Brown K. T., Barott K. L. Coral histology reveals consistent declines in tissue integrity during a marine heatwave despite differences in bleaching severity. PeerJ, 13:e18654, (2005), 10.7717/peerj.18654

7. Lyndby, N. H. et al. The mesoglea buffers the physico-chemical microenvironment of photosymbionts in the upside-down jellyfish Cassiopea sp. 1–12 (2022) doi:10.3389/fevo.2023.1112742.

8. Chan, W. Y. et al. Heat-Evolved Microalgae (Symbiodiniaceae) Are Stable Symbionts and Influence Thermal Tolerance of the Sea Anemone Exaiptasia diaphana. Environ. Microbiol. 27, 1–18 (2025).

9. Currie, A. R. et al. Marine microbial gene abundance and community composition in response to ocean acidification and elevated temperature in two contrasting coastal marine sediments. Front. Microbiol. 8, 1–17 (2017).

10. Zhang, X. H., Ahmad, W., Zhu, X. Y., Chen, J. & Austin, B. Viable but nonculturable bacteria and their resuscitation: implications for cultivating uncultured marine microorganisms. Mar. Life Sci. Technol. 3, 189–203 (2021).

11. Yilmaz, P., Yarza, P., Rapp, J. Z. & Glöckner, F. O. Expanding the world of marine bacterial and archaeal clades. Front. Microbiol. 6, 1–29 (2016).

12. Montaño-Salazar, S., Quintanilla, E. & Sánchez, J. A. Microbial shifts associated to ENSO-derived thermal anomalies reveal coral acclimation at holobiont level. Sci. Rep. 13, 1–11 (2023).

13. Moran M. A., The global ocean microbiome. Science350, aac8455, (2015). DOI:10.1126/science.aac8455

14. Freeman, C. J., Stoner, E. W., Easson, C. G., Matterson, K. O. & Baker, D. M. Symbiont carbon and nitrogen assimilation in the Cassiopea Symbiodinium mutualism. Mar. Ecol. Prog. Ser. 544, 281–286 (2016).

15. Jorissen, H. et al. Coral larval settlement preferences linked to crustose coralline algae with distinct chemical and microbial signatures. Sci. Rep. 11, 1–11 (2021).

16. Weiland-Bräuer, N. et al. The native microbiome is crucial for offspring generation and fitness of aurelia aurita. MBio 11, 1–20 (2020).

17. Murillo-Rincon, A. P. et al. Spontaneous body contractions are modulated by the microbiome of Hydra. Sci. Rep. 7, 1–9 (2017).

18. Krediet, C. J., Ritchie, K. B., Alagely, A. & Teplitski, M. Members of native coral microbiota inhibit glycosidases and thwart colonization of coral mucus by an opportunistic pathogen. ISME J. 7, 980–990 (2013).

19. Neave, M. J., Michell, C. T., Apprill, A. & Voolstra, C. R. Endozoicomonas genomes reveal functional adaptation and plasticity in bacterial strains symbiotically associated with diverse marine hosts. Sci. Rep. 7, 1–12 (2017).

20. Sweet, M. J. & Bulling, M. T. On the importance of the microbiome and pathobiome in coral health and disease. Front. Mar. Sci. 4, 1–11 (2017).

21. Sawiccy V, Tjandra NW, Maruyama S, Ruggeri M, Vo C, Harmon LA, Poole AZ and Weis VM, Regulatory role of NADPH oxidases in symbiosis and dysbiosis in the sea anemone Aiptasia. Front. Mar. Sci. 12:1596098, (2025). doi: 10.3389/fmars.2025.1596098

22. Cui, G. et al. Molecular insights into the Darwin paradox of coral reefs from the sea anemone Aiptasia. Sci. Adv. 9, 1–12 (2023).

23. Randle, J. L., Cárdenas, A., Gegner, H. M., Ziegler, M. & Voolstra, C. R. Salinity-Conveyed Thermotolerance in the Coral Model Aiptasia Is Accompanied by Distinct Changes of the Bacterial Microbiome. Front. Mar. Sci. 7, 1–12 (2020).

24. Pollock, F. J. et al. Coral-associated bacteria demonstrate phylosymbiosis and cophylogeny. Nat. Commun. 9, 1–13 (2018).

25. Huggett, M. J. & Apprill, A. Coral microbiome database : Integration of sequences reveals high diversity and relatedness of coral-associated microbes. Environ. Microbiol. Rep. 11, 372–385 (2019).

26. Sharp, K. H., Pratte, Z. A., Kerwin, A. H., Rotjan, R. D. & Stewart, F. J. Season, but not symbiont state, drives microbiome structure in the temperate coral Astrangia poculata. Microbiome 5, 120 (2017).

27. Curtis, E., Moseley, J., Racicot, R. & Wright, R. M. Bacterial microbiome variation across symbiotic states and clonal lines in a cnidarian model. Front. Mar. Sci. 10, 1–9 (2023).

28. Maire, J., Blackall, L. L. & van Oppen, M. J. H. Microbiome characterization of defensive tissues in the model anemone Exaiptasia diaphana. BMC Microbiol. 21, 1–12 (2021).

29. Brown, T., Otero, C., Grajales, A., Rodriguez, E. & Rodriguez-Lanetty, M. Worldwide exploration of the microbiome harbored by the cnidarian model, Exaiptasia pallida (Agassiz in Verrill, 1864) indicates a lack of bacterial association specificity at a lower taxonomic rank. PeerJ 2017, (2017).

30. Baldassarre, L., Ying, H., Reitzel, A. M., Franzenburg, S. & Fraune, S. Microbiota mediated plasticity promotes thermal adaptation in the sea anemone Nematostella vectensis. Nat. Commun. 13, (2022).

31. Gilbert, J. A., Hill, R., Doblin, M. A. & Ralph, P. J. Microbial consortia increase thermal tolerance of corals. Mar. Biol. 159, 1763–1771 (2012).

32. Ahmed, H. I., Herrera, M., Liew, Y. J. & Aranda, M. Long-term temperature stress in the Coral Model Aiptasia supports the ‘anna Karenina principle’ for bacterial microbiomes. Front. Microbiol. 10, 1–11 (2019).

33. Tignat-Perrier, R., van de Water, J. A. J. M., Allemand, D. & Ferrier-Pagès, C. Holobiont responses of mesophotic precious red coral Corallium rubrum to thermal anomalies. Environ. Microbiome 18, 1–14 (2023).

34. Masson-Delmotte, V. et al. IPCC 2021: Summary for Policy Makers. Cambridge University Press, (2021). doi:10.1017/9781009157896.001.3.

35. McDevitt-Irwin, J. M., Baum, J. K., Garren, M. & Vega Thurber, R. L. Responses of coral-associated bacterial communities to local and global stressors. Front. Mar. Sci. 4, 1–16 (2017).

36. Carabantes, N., Cerqueda-García, D., García-Maldonado, J. Q. & Thomé, P. E. Changes in the Bacterial Community Associated With Experimental Symbiont Loss in the Mucus Layer of Cassiopea xamachana Jellyfish. Front. Mar. Sci. 9, 1–13 (2022).

37. Bollati, E., Hughes, D. J., Suggett, D. J., Raina, J.-B. & Kühl, M. Microscale sampling of the coral gastric cavity reveals a gut-like microbial community. bioRxiv 2024.05.20.594925 (2024).

38. Yamano, R. et al. Genome taxonomy of the genus Thalassotalea and proposal of Thalassotalea hakodatensis sp.nov. isolated from sea cucumber larvae. PLoS One 18, 6–18 (2023).

39. Kim, M., Cha, I. T., Lee, K. E., Lee, E. Y. & Park, S. J. Genomics reveals the metabolic potential and functions in the redistribution of dissolved organic matter in marine environments of the genus thalassotalea. Microorganisms 8, 1–17 (2020).

40. Gerhart, J. G., Moses, A. S. & Raghavan, R. A Francisella-like endosymbiont in the Gulf Coast tick evolved from a mammalian pathogen. Sci. Rep. 6, 1–6 (2016).

41. Schrallhammer, M., Schweikert, M., Vallesi, A. et al. Detection of a Novel Subspecies of Francisella noatunensis as Endosymbiont of the Ciliate Euplotes raikovi. Microb Ecol 61, 455–464 (2011). 10.1007/s00248-010-9772-9

42. Colquhoun, D. J. & Duodu, S. Colquhoun etal francisella 16srRNA.pdf. Vet. Resea 1–15 (2011).

43. Silva-Lima, A. W. et al. Mussismilia braziliensis White Plague Disease Is Characterized by an Affected Coral Immune System and Dysbiosis. Microb. Ecol. 81, 795–806 (2021).

44. Davison, H. R., Hurst, G. D. D. & Siozios, S. ‘Candidatus Megaira’ are diverse symbionts of algae and ciliates with the potential for defensive symbiosis. Microb. Genomics 9, (2023).

45. Klinges, J. G. et al. Phylogenetic, genomic, and biogeographic characterization of a novel and ubiquitous marine invertebrate-associated Rickettsiales parasite, Candidatus Aquarickettsia rohweri, gen. nov., sp. nov. ISME J. 13, 2938–2953 (2019).

46. Pasqualetti, C., Szokoli, F., Rindi, L., Petroni, G. & Schrallhammer, M. The Obligate Symbiont “Candidatus Megaira polyxenophila” Has Variable Effects on the Growth of Different Host Species. Front. Microbiol. 11, 1–10 (2020).

47. Ben-Haim, Y. et al. Vibrio coralliilyticus sp. nov., a temperature-dependent pathogen of the coral Pocillopora damicornis. Int. J. Syst. Evol. Microbiol. 53, 309–315 (2003).

48. Saad, O. S. et al. Genome Size, rDNA Copy, and qPCR Assays for Symbiodiniaceae. Front. Microbiol. 11, 1–16 (2020).

49. Hartman, L. M., Blackall, L. L. & van Oppen, M. J. H. Antibiotics reduce bacterial load in Exaiptasia diaphana, but biofilms hinder its development as a gnotobiotic coral model. Access Microbiol. 4, (2022).

50. Simpson, G. L. gratia: An R package for exploring generalized additive models. J. Open Source Softw. 9, 6962 (2024).

51. Parada, A. E., Needham, D. M. & Fuhrman, J. A. Every base matters: Assessing small subunit rRNA primers for marine microbiomes with mock communities, time series and global field samples. Environ. Microbiol. 18, 1403–1414 (2016).

52. Apprill, A., Mcnally, S., Parsons, R. & Weber, L. Minor revision to V4 region SSU rRNA 806R gene primer greatly increases detection of SAR11 bacterioplankton. Aquat. Microb. Ecol. 75, 129–137 (2015).

53. Callahan, B. J. et al. DADA2: High-resolution sample inference from Illumina amplicon data. Nat. Methods 13, 581–583 (2016).

54. R-Core-Team (Foundation for Statistical Computing). R: A Language and Environment for Statistical Computing. at https://www.r-project.org/ (2022).

55. Quast, C. et al. The SILVA ribosomal RNA gene database project: Improved data processing and web-based tools. Nucleic Acids Res. 41, 590–596 (2013).

56. McMurdie, P. J. & Holmes, S. Phyloseq: An R Package for Reproducible Interactive Analysis and Graphics of Microbiome Census Data. PLoS One 8, (2013).

57. Davis, N. M., Proctor, Di. M., Holmes, S. P., Relman, D. A. & Callahan, B. J. Simple statistical identification and removal of contaminant sequences in marker-gene and metagenomics data. Microbiome 6, 1–14 (2018).

58. Oksanen, J. et al. Vegan: Community Ecology Package, URL: https://cran.r-project.org/web/packages/vegan (2024).

59. Barnett, D., Arts, I. & Penders, J. microViz: an R package for microbiome data visualization and statistics. J. Open Source Softw. 6, 3201 (2021).

60. Martinez Arbizu, P. pairwiseAdonis: Pairwise Multilevel Comparison using Adonis. at https://github.com/pmartinezarbizu/pairwiseAdonis (2017).

61. Lahti, L. & Shetty, S. Microbiome R package, URL: https://microbiome.github.io/tutorials/ (2019).

62. Dörr, M. et al. Short-term heat stress assays resolve effects of host strain, repeat stress, and bacterial inoculation on Aiptasia thermal tolerance phenotypes. Coral Reefs 42, 1271–1281 (2023).

63. Shaffer, M., Thurimella, K., Sterrett, J. D. & Lozupone, C. A. SCNIC: Sparse correlation network investigation for compositional data. Mol. Ecol. Resour. 23, 312–325 (2023).

